# The essential role of sodium bioenergetics and ATP homeostasis in the developmental transitions of a cyanobacterium

**DOI:** 10.1101/2020.12.11.421602

**Authors:** Sofia Doello, Markus Burkhardt, Karl Forchhammer

## Abstract

The ability to resume growth after a dormant period is an important strategy for the survival and spreading of bacterial populations. Energy homeostasis is critical in the transition into and out of a quiescent state. Synechocystis sp. PCC 6803, a non-diazotrophic cyanobacterium, enters metabolic dormancy as a response to nitrogen starvation. We used Synechocystis as a model to investigate the regulation of ATP homeostasis during dormancy and unraveled a critical role for sodium bioenergetics in dormant cells. During nitrogen starvation, cells reduce their ATP levels and engage sodium bioenergetics to maintain the minimum ATP content required for viability. When nitrogen becomes available, energy requirements rise, and cells immediately increase ATP levels employing sodium bioenergetics and glycogen catabolism. These processes allow them to restore the photosynthetic machinery and resume photoautotrophic growth. Our work reveals a precise regulation of the energy metabolism essential for bacterial survival during periods of nutrient deprivation.

## Introduction

Dormant microorganisms are vastly represented in natural environments (Greening et al., 2019). Dormancy highly contributes to the survival of bacterial populations, the spreading of bacterial pathogens and the development of antibiotic resistances (Lewis, 2010). The molecular processes that lead bacterial cells into a dormant state are very diverse, but are generally characterized by growth arrest and residual metabolic activity (Rittershaus et al., 2013). Despite having a reduced metabolism, dormant cells still require energy for maintenance (Greening et al., 2019). In fact, energy homeostasis is known to be critical for the survival of dormant cells (Rittershaus et al., 2013). However, how the energy metabolism is regulated when bacterial cells enter and exit periods of dormancy is poorly understood due to the lack of appropriate model systems.

Cyanobacteria represent a highly diverse group of prokaryotes endowed with the ability to adapt to changing environmental conditions. This feature has allowed them to colonize a wide range of ecosystems (Houmard, 1995). One of the most common hurdles cyanobacterial cells face in natural environments is limitation of a combined nitrogen source (Vitousek & Howarth, 1990). *Synechocystis* sp. PCC 6803 (hereafter *Synechocystis*) is a non-diazotrophic cyanobacterial strain that survives periods of nitrogen starvation by entering metabolic quiescence, thus representing a good model to study fundamental aspects of bacterial dormancy (Klotz et al., 2016). *Synechocystis* can survive prolonged periods of nitrogen starvation by undergoing nitrogen-chlorosis, a process characterized by the degradation of most of the thylakoid membranes. Cells enter cell cycle arrest and shut down their metabolism. Most of the photosynthetic apparatus is degraded, leaving cells with only residual photosynthetic capacity, and energetically costly processes, like anabolic reactions, are halted (Klotz et al., 2016). In this resting state, the intracellular ATP concentration is about ¼ of the level during vegetative growth (Doello et al., 2018). In addition, as cells are degrading most of their cellular components, they synthesize reserve polymers, which are essential for exiting dormancy and resuming growth. Glycogen has been described as the main storage molecule during nitrogen starvation: Its synthesis and degradation are crucial for cell survival under these conditions (Doello et al., 2018; Klotz et al., 2016; Klotz & Forchhammer, 2017).

When nitrogen becomes available to cells in nitrogen-chlorosis, they immediately initiate a highly organized resuscitation program, which has been overall well characterized (Klotz et al., 2016; Spät et al., 2018). During the first stages of the resuscitation process, cells catabolize the accumulated glycogen to obtain the necessary energy and metabolic intermediates to restore all cellular components that had been degraded during chlorosis. When the photosynthetic machinery is restored, cells switch back to phototrophic metabolism (Klotz et al., 2016). Upon nitrogen addition, the energy requirement of chlorotic cells suddenly increases due to the initiation of energy consuming anabolic reactions, such as the glutamine synthetase/glutamate synthase (GS/GOGAT) reaction cycle. Concomitantly with the increased energy demand, the low intracellular ATP concentration of dormant cells rapidly increases to an intermediate level, which represents approximately 50% of the ATP content of a vegetative growing cell (Doello et al., 2018). So far, how dormant cells produce this ATP has remained unknown. Intriguingly, we observed a rapid increase in ATP levels also in mutant cells unable to degrade glycogen (Doello et al., 2018). This observation prompted us to investigate the source of the rise in the cellular ATP content in cells that initiate the resuscitation program. The aim of this study was to reveal how dormant cells maintain the required ATP levels to keep viability, and how they obtain the necessary energy to awaken from dormancy.

## Results

### The rapid ATP increase in response to nitrogen availability is independent of glycogen degradation and photosynthesis

Upon nitrogen addition, cells start the resuscitation program and their energy demands become higher. Cells must then begin producing ATP to support nitrogen assimilation and biosynthetic processes. We measured the intracellular ATP content within the first hour of resuscitation and found that 20 minutes after the addition of sodium nitrate an increase of ~ 50 % in the amount of ATP is observed, and these levels are maintained for the first hour of recovery (**Figure 1**). This ATP increase constitutes the fastest measured response of chlorotic cells to the presence of nitrogen (Klotz et al., 2016), but how cells produce it or what induces its synthesis is not yet understood.

**Figure 1.**
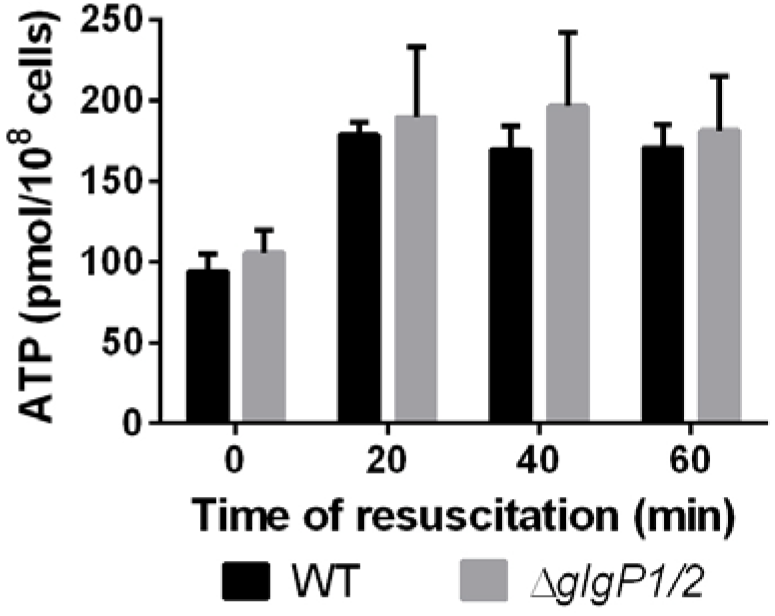
The rapid increase in ATP levels upon sodium nitrate addition is independent of glycogen respiration. ATP content normalized to 1 x 10^8^ cells of WT and Δ*glgP1/2* chlorotic cells after addition of 17 mM NaNO_3_. At least three biological replicates were measured; error bars represent the SD.

A rise in the ATP levels might seem an obvious consequence of the activation of a metabolism that was dormant and enters a transient phase of heterotrophy. Therefore, it is tempting to assume that the increased ATP values at the start of resuscitation come from the onset of glycogen catabolism, which is induced soon after the addition of sodium nitrate to dormant cells (Klotz et al., 2016). Previously, we observed that a mutant lacking the glycogen phosphorylases (Δ*glgP1/2*) displayed elevated ATP levels 3 hours after sodium nitrate addition, but these cells did not further recover. Here, we compared the short-term response in ATP levels between Δ*glgP1/2* and wild-type (WT) in more detail and found that sodium nitrate triggered a similar rapid ATP increase in the Δ*glgP1/2* mutant as it did in the WT, implying that the rapid onset of ATP synthesis does not depend on glycogen catabolism (**Figure 1**). Respiration of other metabolites can be excluded, since Δ*glgP1/2* does not show any oxygen consumption upon addition of sodium nitrate. In fact, the rise in ATP levels happened before cells perform respiration at full capacity. During nitrogen-chlorosis, cells display a residual photosynthetic activity, which is completely repressed after a few hours of resuscitation, when respiration and degradation of glycogen are fully operating (Doello et al., 2018). Pulse-amplitude modulation (PAM) fluorometry measurements revealed that 1 hour after nitrate addition, much after an ATP increase is measurable, glycogen catabolism has not yet suppressed PSII activity (**Figure S1**). After 2 hours of resuscitation, when cells are fully respirating, the PSII activity disappears and only resumes when cells have partially restored their photosynthetic machinery (~ 12 h after nitrate addition). Thus, the observed increase in ATP levels during early resuscitation could depend on photosynthesis instead of respiration. To test this possibility, the ATP content of cells that had been incubated in the dark was measured (**Figure 2A**). Although ATP levels were overall lower in the dark than in the light, addition of sodium nitrate caused a similar increase under both conditions, indicating that photosynthesis is not responsible for the rapid ATP increase after addition of sodium nitrate. To completely exclude the role of photosynthesis on the rise of ATP levels, we treated chlorotic cells with different photosynthetic inhibitors. Exposure to dichlorophenyl dimethylurea (DCMU), which blocks the electron transfer from PSII to the plastoquinone (PQ) (**Figure 2B**), dibromthymonchion (DBMIB), which inhibits the electron flow from PQ to the cytochrome b_6_f complex (Cyt b_6_f), and Antimycin A, which disrupts the Q cycle in Cyt b_6_f (**Figure 2C**) did not affect the cell’s ability to produce ATP after addition of sodium nitrate.

**Figure 2.**
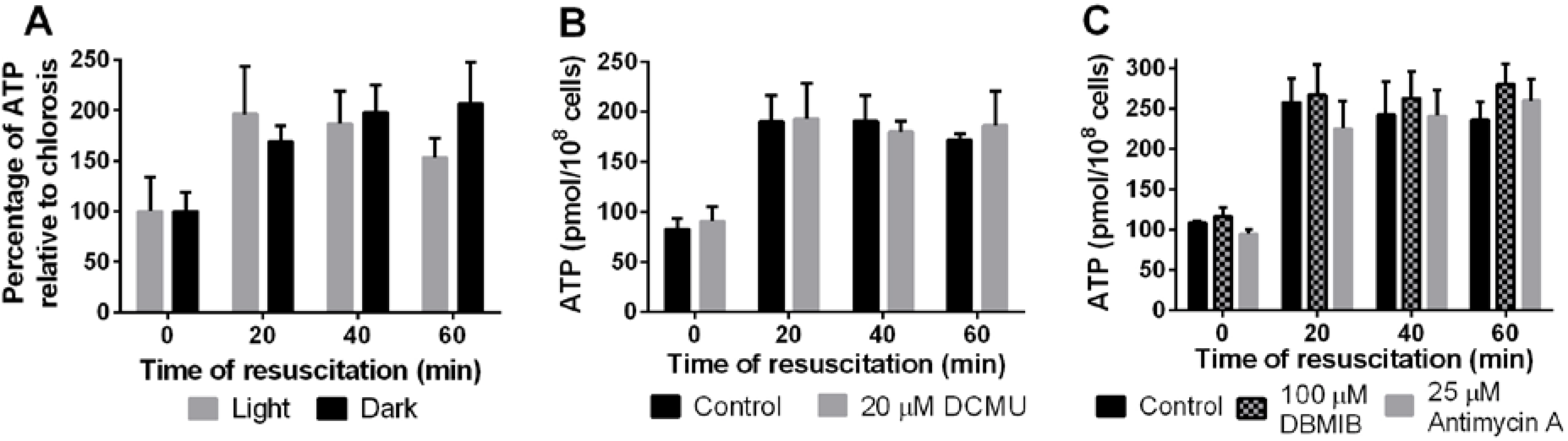
The rapid increase in ATP levels upon sodium nitrate addition is independent of photosynthesis. ATP content normalized to 1 x 10^8^ cells of WT chlorotic cells treated with (A) WT chlorotic cells after incubation for 1 h in darkness and addition of 17 mM NaNO_3_. (B) 20 μM DCMU and (C) 100 μM DBMIB and 25 μM Antimycin A. Cells were treated for 5 min before the first measurement (0 min). Resuscitation was then induced by addition of 17 mM NaNO_3_. At least three biological replicates were measured; error bars represent the SD.

### The ATP increase relies on a sodium-motive force

Respiration and photosynthesis are the two main bioenergetic processes that generate an electrochemical proton gradient that can be used by the ATP synthase to power ATP production. When both processes were blocked, nitrogen-starved cells could still increase ATP levels upon addition of sodium nitrate. To elucidate the contribution of proton-motive force on the rise in ATP levels, chlorotic cells were treated with the protonophores carbonyl cyanide m-chlorophenyl hydrazone (CCCP) and 2,3-dinitrophenol (DNP). Protonophores make membranes permeable to protons, thus destroying proton gradients. Intriguingly, treatment with CCCP and DNP did not abolish the rise in ATP levels (**Figure 3**), indicating that ATP synthesis does not depend on an electrochemical proton gradient. However, protons are not the only ion motive force that can be coupled to ATP synthesis, as some ATP synthases can also use a sodium-motive force to power ATP production (Schulz et al., 2013). Sodium ions are more abundant in the extracellular medium than in the cytoplasm, thus forming a gradient across the plasma membrane that can be utilized by sodium-binding ATP synthases to produce ATP. Besides the thylakoidal ATP synthases, which translocate the protons accumulated in the thylakoid lumen into the cytoplasm to produce ATP, *Synechocystis* also possesses ATP synthases in the plasma membrane (Huang et al., 2002), which might use a sodium-motive force.

**Figure 3.**
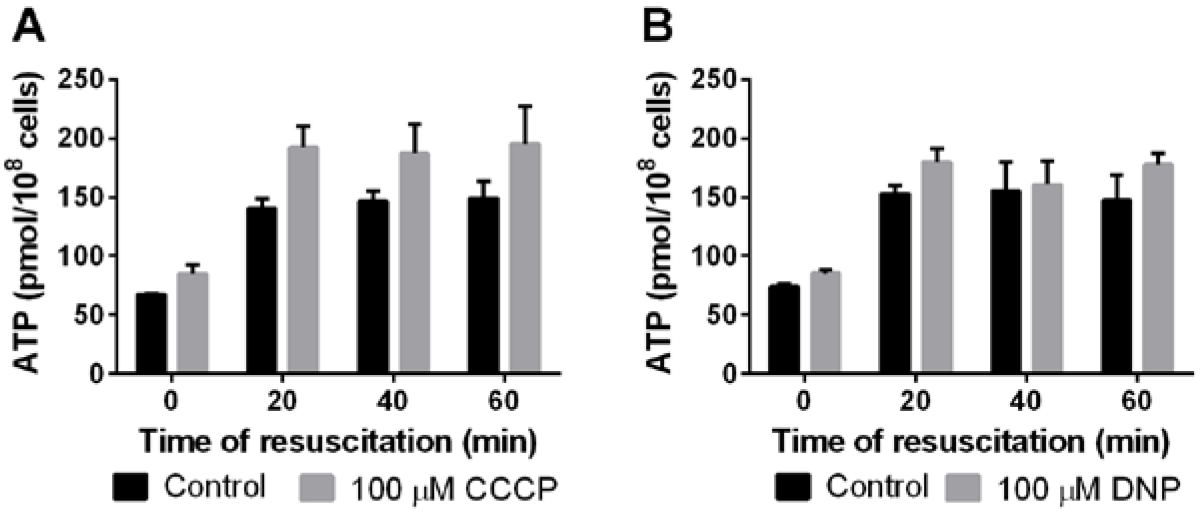
Dissipation of the proton gradient does not inhibit the rapid increase in ATP levels. ATP content normalized to 1 x 10^8^ cells of WT chlorotic cells treated with (A) 100 μM CCCP and (B) 100 μM DNP. Cells were treated for 5 min before the first measurement (0 min). Resuscitation was then induced by addition of 17 mM NaNO_3_. At least three biological replicates were measured; error bars represent the SD.

The above experiments were performed by adding 17 mM sodium nitrate to chlorotic cells in nitrogen-free BG_11-0_ medium, increasing the sodium concentration 4-fold. This raised the question whether the rapid increase of intracellular ATP is connected to the sudden rise in sodium levels. To test this, recovery experiments were performed by the addition of 17 mM potassium nitrate or 5 mM ammonium chloride to cells in BG_11-0_ (**Figure 4A**). In these cases, the concentration of sodium remained constant. Remarkably, the ATP increase was significantly lower than in the previous experiments with the addition of 17 mM sodium nitrate. The rapid rise in ATP levels could be restored when 17 mM sodium chloride was added together with potassium nitrate or ammonium chloride to dormant cells (**Figure 4B**). When sodium was completely removed from the medium by repeated washing with BG_11-0-Na_ (in which all sodium salts have been substituted by potassium salts), addition of potassium nitrate triggered almost no increase of ATP levels (**Figure 4C**). These results demonstrated that sodium plays an important role in ATP synthesis in chlorotic cells. However, whether or not the addition of a nitrogen source also contributes to the rise in ATP levels remained unclear, since cells in sodium-free medium still reacted to the addition of potassium nitrate with a small increase in the concentration of ATP. To address this question, sodium and nitrogen were added to dormant cells sequentially. As shown in **Figure 4D**, the sole addition of 17 mM sodium chloride to chlorotic cells in BG_11-0_ caused a partial increase of the ATP levels within 20 minutes, compared to the standard resuscitation experiment. When 20 minutes after supplementation with sodium chloride a nitrogen source was added to the cells, either as potassium nitrate (**Figure 4D, column A**) or as ammonium chloride (**Figure 4D, column B**), a further rise in ATP levels was observed after 1 hour. This indicates that the rise in ATP levels that was initially observed when sodium nitrate was added to chlorotic cells has two components: One due to the increase in the sodium concentration, and another one due to the presence of a nitrogen source. To distinguish whether the cells directly sense the nitrogen source or detect it through initiating assimilation via the GS-GOGAT cycle, cells were treated with the GS inhibitor L-methionine sulfoximine (MSX). Indeed, this treatment completely abolished the nitrogen-dependent component of the ATP increase (**Figure 4D, column C**), indicating that cells respond to the assimilation of ammonium rather than to the external presence of a combined nitrogen source. Interestingly, the Δ*glgP1/2* mutant only reacted to sodium and did not show the nitrogen-dependent component of the ATP increase (**Figure 4D, column D**). These results show that both, activation of nitrogen assimilation and glycogen degradation, are required for the nitrogendependent ATP increase.

**Figure 4.**
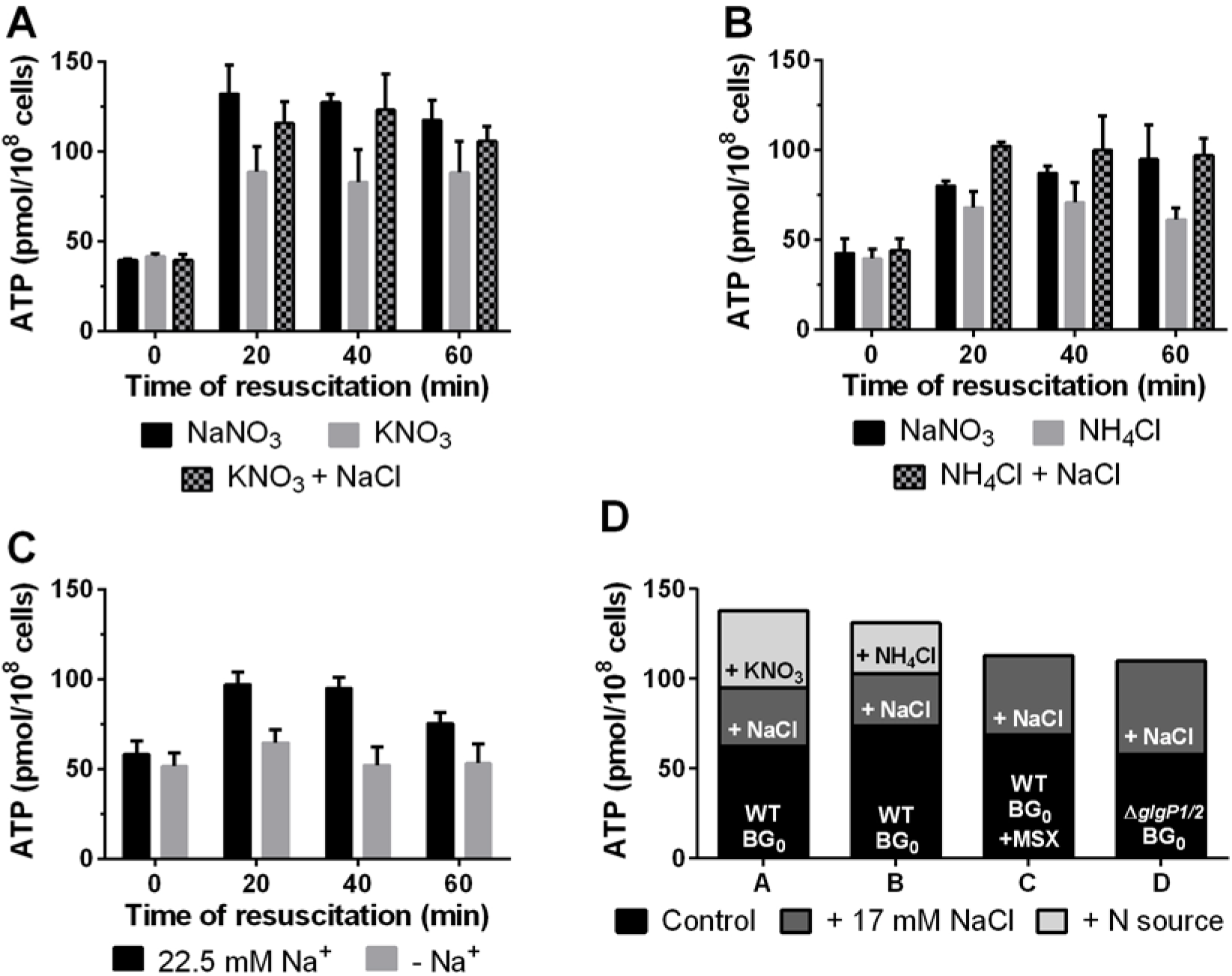
The rise of cellular ATP upon sodium nitrate addition is a response to both, increased sodium concentrations and nitrogen assimilation. ATP content normalized to 1 x 10^8^ cells of WT chlorotic cells. (A) Cells were resuscitated using either 17 mM NaNO_3_, 17 mM KNO_3_, or 17 mM KNO_3_ + 17 mM NaCl. (B) Cells were resuscitated using either 17 mM NaNO_3_, 5 mM NH_4_Cl, or 5 mM NH_4_Cl + 17 mM NaCl. (C) Cells were washed twice with BG_11-0-Na_ sodium-free medium and resuscitated with 17 mM KNO_3_. (D) ATP content normalized to 1 x 10^8^ cells of chlorotic cells in BG_11-0_ (black), after supplementation with 17 mM NaCl (dark grey) and after additional supplementation with a nitrogen source (light grey). Column A: untreated WT chlorotic cells supplemented with 17 mM NaCl and 17 mM KNO_3_. Column B: untreated WT chlorotic cells supplemented with 17 mm NaCl and 5 mM NH_4_Cl. Column C: WT chlorotic cells treated with 200 μM MSX and supplemented with 17 mM NaCl and 17 mM KNO_3_. Column D: untreated Δ*glgP1/2* chlorotic cells supplemented with 17 mM NaCl and 17 mM KNO_3_. At least three biological replicates were measured; error bars represent the SD.

In order to corroborate the role of sodium in ATP synthesis during chlorosis, nitrogen-starved cells were treated with monensin, a sodium ionophore, and ethyl-isopropyl amiloride (EIPA), an inhibitor of sodium channels and sodium/proton antiport. To exclude any indirect effects caused by possible interference of the inhibitors with nitrate transport, the effect of monensin and EIPA on the ATP content was measured after adding a combination of ammonium chloride and sodium chloride to chlorotic cells. Treatment with monensin led to lower ATP levels than the untreated control (**Figure 5A**). More strikingly, exposure to EIPA completely abolished the increase in ATP levels (**Figure 5B**), proving the key role of sodium in the bioenergetics of chlorotic cells.

**Figure 5.**
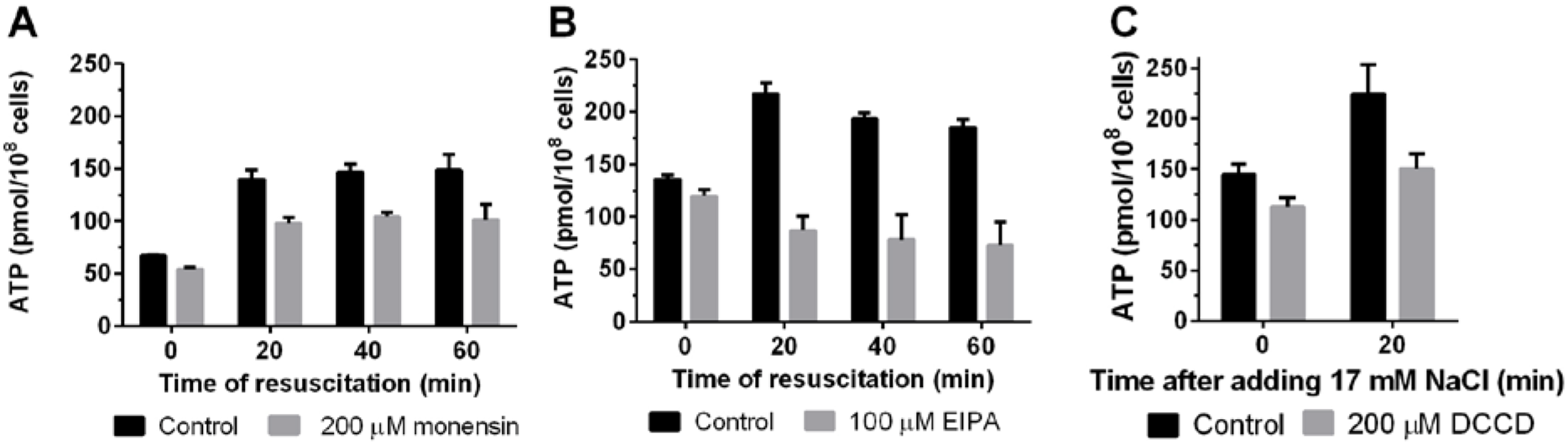
Dissipation of the sodium ion gradient, inhibition of sodium ion transport across the membrane, and inhibition of F-ATPases dampens the rapid rise in intracellular ATP. ATP content normalized to 1 x 10^8^ cells of WT chlorotic cells treated with (A) 200 μM monensin, (B) 100 μM EIPA, and (C) 200 μM DCCD. Cells were treated for 5 min before the first measurement (0 min). In (A) and (B), resuscitation was induced by addition of 5 mM NH_4_Cl + 17 mM NaCl. In (C) only 17 mM NaCl was added to induce sodium-dependent ATP synthesis. At least three biological replicates were measured; error bars represent the SD.

To ascertain whether the ATP synthases are responsible for the sodium-dependent component of the ATP increase, chlorotic cells were treated with the potent F-ATPase inhibitor N, N’-dicyclohexylcarbodiimide (DCCD). Cells that were exposed to DCCD showed a reduced response to the addition of sodium chloride as compared to untreated cells (**Figure 5C**), confirming that the sodium-dependent ATP increase relies on the activity of the ATP synthases.

### Sodium requirement depends on the cellular growth stage

Hitherto, it remained unclear if cells engage sodium bioenergetics exclusively during nitrogen-chlorosis or if sodium-dependent ATP synthesis is a part of *Synechocystis* metabolism in general. To answer this question, vegetative cells were treated with monensin and EIPA. The ATP content of vegetative cells was not affected after treatment with monensin for 30 minutes (**Figure 6A**). By contrast, EIPA slightly reduced the ATP levels by 25% in vegetative cells (**Figure 6B**). However, treatment with EIPA also completely inhibited PSII activity (**Figure S2**), suggesting that the observed lower ATP levels might be a consequence of the inhibitory effect of EIPA on photosynthesis rather than a direct effect on sodium-dependent ATP synthesis. These results indicate that, while sodium bioenergetics plays a key role during nitrogen starvation, vegetative cells do not rely on sodium-dependent ATP synthesis.

**Figure 6.**
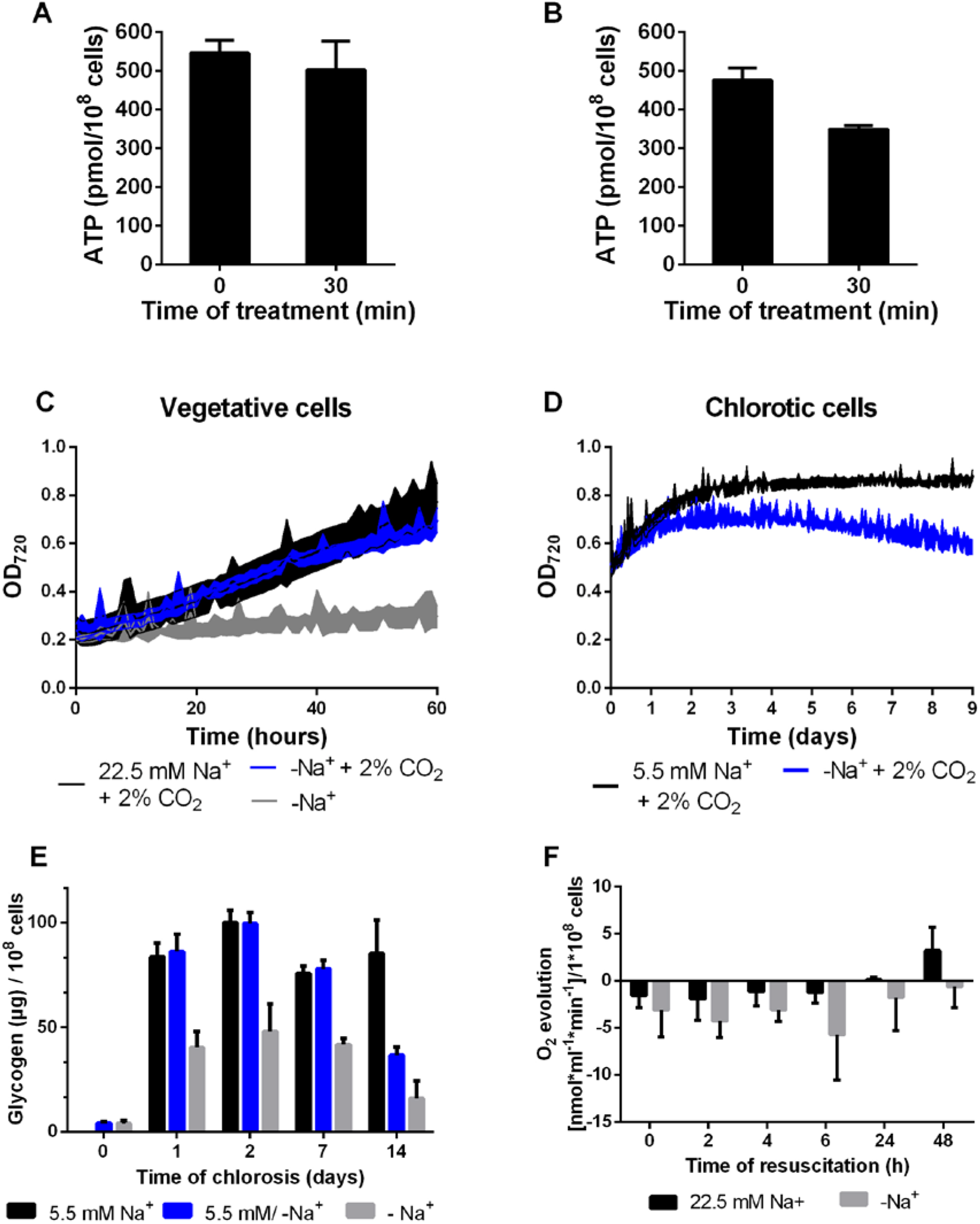
Sodium is required for bicarbonate uptake but not for ATP synthesis during vegetative growth. (A) ATP content normalized to 1 x 10^8^ cells of vegetative cells treated with 200 μM monensin for 30 minutes. (B) ATP content normalized to 1 x 10^8^ cells of vegetative cells treated with 100 μM EIPA for 30 minutes. (C) Optical density at 720 nm of vegetative cells in regular BG_11_ suppleented with 2 % CO_2_ (black line) and in BG_11-Na_ sodium-free medium with ambient air (grey line) and 2% CO_2_ supplementation (blue line). (D) Optical density at 720 nm of chlorotic cells in regular BG_11-0_ supplemented with 2 % CO_2_ (black line) and in BG_11-0-Na_ sodium-free medium supplemented with 2% CO_2_ (blue line). (E) Glycogen content normalized to 1 x 10^8^ cells throughout chlorosis of WT cells in medium containing 5.5 mM sodium (black bars, standard conditions), sodium-free medium (grey bars) and cells that were cultivated in standard conditions, i.e. 5.5 mM sodium, for 24 h and then transferred to sodium-free medium (blue bars). (F) Oxygen evolution of resuscitating WT cells in medium containing 22.5 mM sodium (black bars, standard conditions) and sodium-free medium (grey bars). At least three biological replicates were measured; error bars represent the SD.

To further elucidate the role of sodium on the metabolism of *Synechocystis*, vegetative and nitrogen-starved cells were cultivated in sodium-free medium. Under atmospheric gas conditions in shaking flasks, vegetative cells could not grow in the absence of sodium; however, growth in sodium-free medium could be restored when cells were supplemented with 2% CO_2_ (**Figure 6C**). These findings are explained by the fact that sodium is required for bicarbonate uptake through the SbtA and BicA transporters (Burnap et al., 2015; Shibata et al., 2002). Thus, sodium-dependent bicarbonate transport is essential for growth under conditions of atmospheric CO_2_ supply, but cells do not require sodium with elevated CO_2_ concentrations. Conversely, nitrogen-starved cells showed a decreasing optical density when cultivated in sodium-free medium, even when they were supplemented with 2 % CO_2_ (**Figure 6D**) indicating a requirement for sodium beyond the need for inorganic carbon transport.

In the presence of sodium, during the first 24 hours after nitrogen deprivation, cells synthesize large amounts of glycogen. In sodium-free medium and under atmospheric gas conditions, cells accumulate only ~ 50 % of the amount of glycogen after 2 days of nitrogen starvation as compared to the standard medium, and upon further incubation, glycogen levels further decreased (**Figure 6E**). When cells were nitrogen-starved under standard conditions for 24 hours, until they reached the maximum glycogen content, and were then transferred to sodium-free medium, the glycogen concentration progressively decreased after sodium removal (**Figure 6E**). This finding suggests that the absence of sodium triggers glycogen catabolism. When resuscitation of chlorotic cells in sodium-free medium was initiated by the addition of potassium nitrate (conditions in which only a low ATP increase was observed, see above), they showed higher respiration rates than cells resuscitating under standard conditions (**Figure 6F**). However, these cells were unable to complete the resuscitation process despite being able to switch on the initial steps of the resuscitation metabolism: They never re-greened and eventually lost viability, as shown by the complete loss of photosynthetic activity (**Figure S1**). This further emphasizes the dependence of chlorotic cells on sodium bioenergetics.

### ATP levels are rapidly tuned depending on the metabolic requirements

So far, the analysis of sodium requirement in vegetative and chlorotic cells showed that vegetative cells require sodium for bicarbonate transport, whereas chlorotic cells require sodium for bioenergetics. When vegetative cells are shifted to nitrogen-deprived conditions, they are initially photosynthetically competent. In order to elucidate how ATP levels are affected after transferring vegetative cells to nitrogen-deprived conditions at different sodium concentrations, we analyzed the ATP content of vegetative cells after they were transferred either into regular BG_11-0_ (5.5 mM sodium) or into BG_11-0_ supplemented with 17 mM sodium chloride, which equals the concentration of sodium in BG_11_ (22.5 mM). Already 30 min after shifting to nitrogen-deficient medium, the ATP levels dropped to approximately one third of the initial value, regardless of the sodium concentration. Subsequently, the ATP content was then maintained at this low level during long-term chlorosis (**Figure 7A**). To ensure that the decrease in ATP levels was not simply due to a globally reduced level of nucleotides, we determined the ATP to ADP ratio, which should stay constant in the case of a general adenine nucleotide decrease. As shown in **Figure 7B**, the ratio dropped in a similar manner than ATP levels decreased, indicating a reduced energy charge rather than a decrease of nitrogen-containing compounds after nitrogen removal. This implies that cells specifically adjust ATP levels as a response to severe metabolic imbalance for the consequent need to globally modify cellular processes.

**Figure 7.**
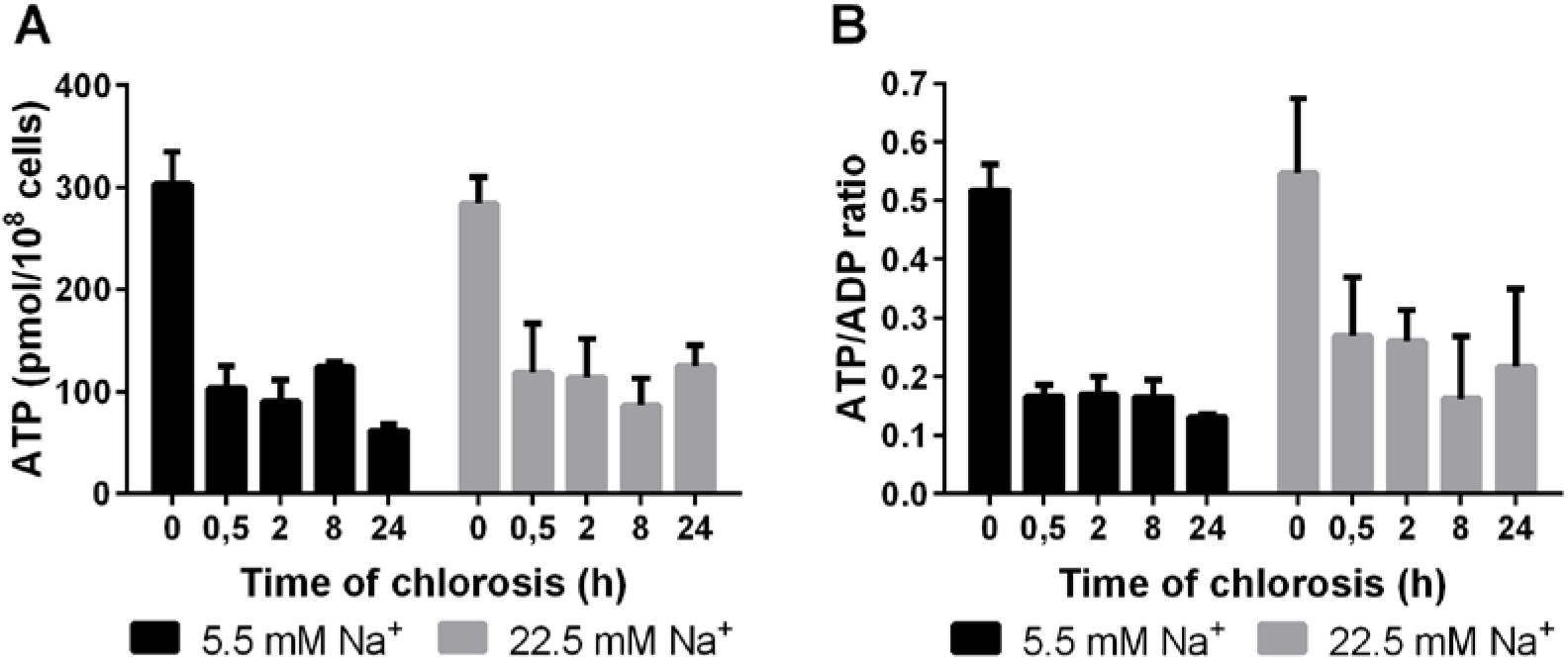
The ATP concentration is rapidly reduced after nitrogen step-down even in the presence of high sodium. (A) ATP content normalized to 1 x 10^8^ cells of WT cells after nitrogen deprivation in standard conditions (5.5 mM sodium, black bars) and high-sodium conditions (22.5 mM sodium, grey bars). (B) ATP/ADP ratio of WT cells after nitrogen deprivation in standard conditions (5.5 mM sodium, black bars) and high-sodium conditions (22.5 mM sodium, grey bars) At least three biological replicates were measured; error bars represent the SD.

## Discussion

### *Synechocystis* engages sodium bioenergetics during nitrogen starvation

During nitrogen-chlorosis, *Synechocystis* re-arranges its metabolism to reach a state of dormancy that allows cell survival for a prolonged starvation time. This metabolic adaptation includes reduction of both energy consumption and production. Thus, chlorotic cells keep ATP low, just at the minimum level to ensure cell survival (~ 50-100 pmol/ 10^8^ cells) (Doello et al., 2018). When a nitrogen source is added to chlorotic cells, the ATP demand increases immediately due to the ammonium-assimilating GS-reaction, which consumes one ATP per ammonium, the concomitant GOGAT reaction and all the following anabolic processes that are induced at the onset of resuscitation (Klotz et al., 2016; Spät et al., 2018). Cells respond accordingly and almost immediately increase their ATP levels by about ~50 % to power the anabolic reactions.

Most of the cellular ATP is produced by the ATP-synthases from ADP and inorganic phosphate. In cyanobacteria, this reaction typically requires an electrochemical proton gradient across the thylakoid membrane, which is generated by photosynthetic or respiratory electron transport (Imashimizu et al., 2011). However, chlorotic cells could still increase ATP levels within several minutes, even when the two main bioenergetic processes that generate a proton gradient were inhibited. We could identify the nature of this increase in the ATP content and dissect it into two components: One that is purely sodium-dependent, and a second one that was triggered by ammonium assimilation and supported by glycogen degradation.

Chlorotic cells have largely degraded their thylakoids, and therefore the space for thylakoidal ATP synthases and proton storage is limited (Klotz et al., 2016). Previous studies have reported the presence of ATP synthases in the plasma membrane of *Synechocystis* (Huang et al., 2002), which suggests that cells could use an extracellular electrochemical gradient to power ATP synthesis. However, cyanobacteria preferably grow under alkaline conditions, where the abundance of protons is extremely low. By contrast, sodium ions are highly abundant in the extracellular medium and cells are exposed to a natural sodium gradient due to sodium extrusion from the cytoplasm. The first component of the rapid rise in intracellular ATP that was triggered by addition of sodium nitrate to chlorotic cells can be explained by an increase in the sodium-motive force. The second component of the ATP increase could be prevented by treatment with MSX, a specific GS inhibitor, and was absent in a mutant unable to degrade glycogen. This suggests that initiation of nitrogen assimilation triggers glycogen catabolism, which supports ATP synthesis via substrate level phosphorylation and, even more efficiently, by supporting respiration. These results show that bioenergetics of chlorotic cells is largely based on sodium, which allows chlorotic cells to keep the minimum intracellular ATP concentration to maintain cell viability during metabolic dormancy, even in an alkaline environment.

### Energy homeostasis in *Synechocystis*

In contrast to chlorotic cells, vegetative cells do not rely on sodium-dependent ATP synthesis. As shown here, and in agreement with previous studies (Burnap et al., 2015; Shibata et al., 2002), vegetative cells require sodium primarily for sodium-dependent bicarbonate uptake. During vegetative growth, the major ATP synthesis machinery is located in the thylakoid membranes, where photosynthetic and respiratory complexes generate a protonmotive force to power ATP synthesis (Imashimizu et al., 2011). Upon nitrogen starvation, nitrogen assimilation and most anabolic processes are halted. Under these circumstances, ATP levels would be expected to increase, since at this point the ATP synthesis machinery is still intact and the most energy consuming reactions in the cell stop taking place. However, when cells were transferred to nitrogen-free medium, a rapid and steep decrease in ATP levels was observed, independently of the concentration of sodium in the medium. The fact that cells respond in this opposite way, decreasing their ATP content instead of increasing it, suggests the existence of a powerful, yet unexplored regulatory mechanism of tuning ATP levels.

Reduced ATP levels have previously been reported in bacterial cells during periods of metabolic dormancy. In *Mycobacterium tuberculosis*, the ATP content in nutrient-starved cells are steadily maintained at a constant level which is 5-fold lower than the levels in growing cells (Rittershaus et al., 2013). However, whether the decreased ATP content is a consequence of a reduced metabolic activity during bacterial dormancy, or if low ATP levels are required to reach this metabolic state, has not been elucidated. In *Synechocystis*, mutants unable to synthesize glycogen (Δ*glg.A1/2* and Δ*glgC*) present higher ATP levels than the WT and fail to perform a proper acclimation response to nitrogen starvation, which leads to death (Gründel *et al*., 2012; Cano *et al*., 2018; Díaz-Troya *et al*., 2020). However, this phenotype is alleviated when synthesis of the osmolyte glucosylglycerol, which is produced from ADP-glucose under conditions of high salt stress, is induced in the Δ*glgA1/2* mutant, showing the importance of an energy dissipation pathway for acclimation to nitrogen starvation (Díaz-Troya et al., 2020). These findings, together with our observation that ATP levels rapidly drop after nitrogen stepdown, even in the presence of high sodium, strongly support the idea that a decreased ATP content is important for adaptation of the metabolism to nitrogen starvation. Reduction of the ATP levels may play a role in re-directing the metabolism into dormancy, since some cellular processes that are important for this transition, such as the formation of protein aggregates, are promoted by decreased cellular ATP concentrations (Pu et al., 2019). Also, ATP has been shown to act as a biological hydrotrope that influences the fluidity of the cytoplasm (Patel et al., 2017). Adaptation of the cytoplasm from a fluid to a glass-like state has important implications on molecular diffusion inside the cells and plays a relevant role in bacterial adaptation to dormancy (Parry et al., 2014). Low ATP levels might be a necessary factor for the transition of the cytoplasm into a glass-like state.

Although deciphering the mechanism that regulates ATP levels when cells are shifted to nitrogen-free medium was not the aim of this study, we observed that glycogen degradation is induced in the absence of sodium in chlorotic cells, probably in an attempt to maintain the cellular ATP content to a minimum level. Similarly, during resuscitation in sodium-free medium, cells respired more (i.e. degraded more glycogen) than in presence of high sodium, most likely to compensate for the lack of sodium-dependent ATP synthesis. These findings support the idea that was already proposed by Cano et al. (Cano et al., 2018). They suggested that glycogen metabolism is controlled by the intracellular energy charge and plays an important role in energy homeostasis in response to the growth phase and the environmental conditions. Nevertheless, the exact molecular mechanism that allows energy dissipation upon nitrogen removal needs yet to be elucidated.

### Proposed mechanism of sodium-dependent ATP synthesis in *Synechocystis*

The ATP synthase is formed by a membrane complex (F_0_), which transports the ions across the membrane, and a cytoplasmic complex (F_1_), where ATP is synthesized. It is in the c-ring within complex F_0_ that ion specificity is determined. In *Synechocystis* this is formed by 14 copies of the subunit c (AtpE) (Pogoryelov et al., 2007; Schulz et al., 2013). Some marine cyanobacteria contain a gene encoding for a sodium-translocating AtpE in addition to the proton-translocating one (Dibrova et al., 2010). This is not the case for *Synechocystis:* In its genome there is just one gene that encodes for AtpE. Whether the c-ring binds protons or sodium ions is not dictated by major structural differences, but by slight variations in the amino acid sequence around the ion-binding site. Both, protons and sodium ions, bind a glutamate residue, and it is the nature of the amino acids surrounding this residue that determine the specificity of the c-ring. Sodium-ATP synthases have several polar groups in their ion-binding site which form a complex network of hydrogen bonds, whereas proton-ATP synthases have more hydrophobic residues (Leone et al., 2015; Schulz et al., 2013). The balance between hydrophobic and polar groups makes the c-rings more or less selective towards one ion or the other. Those ATP synthases that are driven by an electrochemical proton gradient must have a high proton selectivity, since usually the concentration of sodium is much higher than the concentration of protons in physiological conditions. Some ATP synthases, like that from *Methanosarcina acetivorans*, possess proton-specific c-rings, but bind sodium physiologically because their proton specificity is not strong enough to overcome the excess of sodium (Leone et al., 2015; Schlegel et al., 2012). **Figure 8A** shows an alignment of the sequence of *Synechocystis*’ AtpE with those from *Ilyobacter tartaricus, M. acetivorans* and *Arthrospira platensis*, with weak, medium and strong proton selectivity, respectively. There are 5 key amino acids (marked in orange and green) around the glutamate residue where the ion binds (marked in red) that favor sodium-binding. In the c-ring of *I. tartaricus* all of them are present: Sodium ions bind the oxygen atoms in the side chain of E65 (the main ion-binding residue), S66 and Q32, and the backbone carbonyl group of V63; additionally, the sodium ion is stabilized by a water molecule, which interacts with T67 and A64 (**Figure 8B**). *M. acetivorans*’ AtpE possesses those residues that directly interact with the sodium ion, or substitutions that form identical interactions (a serine in the position of S66, a glutamate replacing Q32, which is assumed to be constitutively protonated and form the same interaction than glutamine, and a methionine substituting V63, which only participates in the interaction through the backbone carbonyl group), but lacks those residues that stabilize the water molecule, which gives the c-ring a higher proton selectivity than the one of *I. tartaricus. A. platensis*’ AtpE has an alanine replacing S66 and a leucine in the position of T67; the presence of these hydrophobic residues confers the c-ring high proton selectivity. Around the AtpE ion-binding site, *Synechocystis*’ AtpE has similar key residues to the ones of*M. acetivorans:* It also contains the polar residues that interact with the sodium ion, but not the residues that interact with the stabilizing water molecule. We thus estimated that *Synechocystis* c-ring also presents medium proton selectivity, but can bind sodium if it is in excess and the ATP synthase could therefore be responsible for sodium-dependent ATP synthesis. We could confirm this with the observation that blocking the ATP synthase by treating chlorotic cells with DCCD inhibited the sodium-dependent ATP increase. A moderate proton specificity permits the ATP synthases in the thylakoid membranes to bind protons physiologically, since the concentration of protons in the thylakoid lumen is high. However, those ATP synthases located in the plasma membrane of dormant cells that live in an alkaline environment are more likely to bind sodium. This enzyme promiscuity allows dormant cells to adapt and survive to an environment where the classical ways to obtain energy are limited.

**Figure 8.**
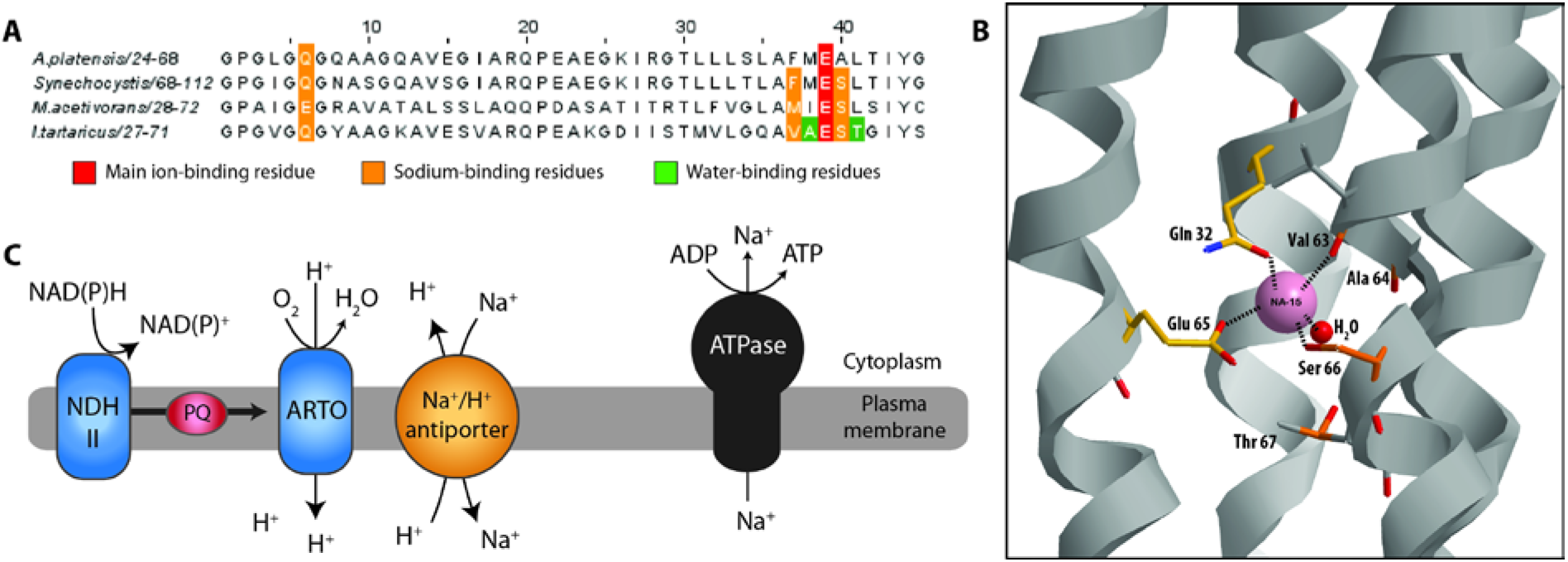
Proposed mechanism of sodium-dependent ATP synthesis in *Synechocystis*. (A) Alignment of the sequence of the ion-binding site of AtpE from *Arthrospira platensis, Synechocystis, Methanosarcina acetivorans* and *Ilyobacter tartaricus*. Residues involved in Na^+^ coordination are indicated in colors. (B) Na^+^ coordination in the c-ring from *I. tartaricus*. (C) Proposed mechanism for maintaining a Na^+^ gradient.

Further efforts will be needed to elucidate how chlorotic cells maintain an electrochemical sodium gradient across the plasma membrane. Previous studies have found evidence of respiratory electron transport in the plasma membrane of *Synechocystis* (Baers et al., 2019; Huang et al., 2002). Baers et al. suggested a simpler electron transport chain for the plasma membrane in which NAD(P)H dehydrogenases type II (NDH II) transfer electrons to the plastoquinone pool (PQ), from where the electrons are further transferred to an alternative respiratory terminal oxidase (ARTO) (Baers et al., 2019). With the currently available data, we propose that the protons transported from the cytoplasm to the periplasmic space by ARTO could be directly used by closely located sodium/protons antiporters to extrude sodium ions from the cytoplasm (**Figure 8C**). In experimental support to this hypothesis, we found that NDH II and sodium/proton antiporters are up-regulated in chlorotic cells (Spät et al., 2018). Interestingly, NdbA (slr0851), one of the three NDH II isoenzymes in *Synechocystis*, is the third most up-regulated protein in chlorotic cells. Moreover, this model is in accordance with the extreme sensitivity of chlorotic cells towards the inhibitor of sodium/proton antiport, EIPA.

Altogether, our study sheds light on the regulation of the energy metabolism during bacterial dormancy, which plays a crucial role in the survival and spread of bacterial populations. It remains to be seen how common the phenomenon of engaging sodium bioenergetics to adjust ATP levels to the specific metabolic requirements of each phase of the life cycle is among bacterial species that undergo similar developmental transitions than *Synechocystis*.

## Acknowledgements

We thank Dr. Libera Lo Presti for her assistance writing this manuscript.

## Authors contribution

S.D. and K.F. designed the experiments. S.D. and M.B. conducted the experiments. S.D. and K.F. analyzed the data and wrote the manuscript.

## Experimental procedures

### Cyanobacterial cultivation conditions

*Synechocystis* WT and Δ*glgP1/2* (Doello et al., 2018) strains were grown in BG_11_ supplemented with 5 mM NaHCO_3_ for vegetative growth, as described previously (Rippka et al., 1979). The concentration of sodium in standard BG_11_ medium is 22.5 mM. Nitrogen starvation was induced as previously described by a 2-step wash with BG_11-0_ medium supplemented with 5 mM NaHCO_3_, which contains all BG_11_ components except for NaNO_3_ (Klotz et al., 2016; Schlebusch & Forchhammer, 2010). The concentration of sodium in standard BG_11-0_ medium is 5.5 mM. Resuscitation was induced by addition of 17 mM NaNO_3_ to cells residing in BG_11-0_ (standard conditions). When indicated, 17 mM NaNO_3_ was substituted by 17 mM KNO_3_ or 5 mM NH_4_Cl in recovery experiments, with or without supplementation with 17 mM NaCl, as specified. When stated, cells were transferred to sodium-free (BG_11-Na_ or BG_11-0-Na_) medium, where all sodium salts were replaced by potassium salts. When specified, cells were treated with the inhibitors DCMU (20 μM), DBMIB (100 μM), Antimycin A (25 μM), CCCP (100 μM), DNP (100 μM), MSX (200 μM), monensin (200 μM), EIPA (100 μM) and DCCD (200 μM) for 5 minutes before the experiment was started unless otherwise indicated. Cultivation was performed with continuous illumination (50 to 60 μmol photons m-2 s-1) and shaking (130 to 140 rpm) at 27 °C. Δ*glgP1/2* pre-cultures were cultivated with the appropriate concentration of antibiotics (Doello et al., 2018). Biological replicates were inoculated with the same pre-cultures, but propagated, nitrogen-starved and resuscitated independently in different flasks under identical conditions.

### Growth curves

Growth curves were generated using a Multi-cultivator OD-1000 with a Gas Mixing System GMS 150 (Photosystems Instruments, Drasov, Czech Republic). Vegetative cells were grown in BG_11_ or BG_11-Na_ medium with and without supplementation with 2% CO_2_. Nitrogen starvation was induced as described above, followed by cultivation in BG_11-0_ or BG_11-0-Na_ medium supplemented with 2 % CO_2_. The OD was monitored at 720 nm. Three biological replicates per condition were measured.

### ATP determination

1 mL aliquots of bacterial cultures were taken and immediately frozen in liquid nitrogen. ATP was extracted by boiling and freezing samples 3 times consecutively (boiling at 100 °C, freezing in liquid nitrogen) and spinning them down at 25,000 g for 1 minute at 4 °C. ATP in the supernatant was quantified with the “ATP determination kit” (Molecular Probes (A22066), Oregon, USA) following the manufacturer’s protocol. 50 μl of a reaction mix containing reaction buffer, luciferin and firefly luciferase were mixed with 10 μl of the samples and the luminescence was quantified in a luminometer (Sirius Luminometer, Berthold Detection Systems). An ATP standard curve was generated and used to calculate ATP content in the collected samples. For every condition, at least three biological replicates were measured.

### ADP determination

1 mL aliquots of bacterial cultures were taken and immediately frozen in liquid nitrogen. ATP was extracted by boiling and freezing samples 3 times consecutively (boiling at 100 °C, freezing in liquid nitrogen) and spinning them down at 25,000 g for 1 minute at 4 °C. ATP in the supernatant was quantified with the “ADP Assay Kit” (MAK133, Sigma-Aldrich, Missouri, USA) following the manufacturer’s protocol. 90 μl of a reaction mix containing reaction buffer, luciferin and firefly luciferase were mixed with 10 μl of the samples and the luminescence was quantified in a luminometer to determine the RLU_ATP_. Subsequently, “ADP enzyme” was added to the samples and the luminescence was measured again after a 2-min incubation to determine the RLU_ADP_. An ADP standard curve was generated. The luminescence corresponding to ADP was calculated (RLU_ADP_-RLU_ATP_) and the ADP content in the samples was determined using the standard curve. For every condition, at least three biological replicates were measured.

### Glycogen determination

Glycogen content was determined as described by Gründel et al. (Gründel et al., 2012) with modifications established by Klotz et al. (Klotz et al., 2016). 2 mL-samples were collected, span down and washed with distilled water. Cells were lysed by incubation in 30% KOH at 95°C for 2h. Glycogen was precipitated by addition of cold ethanol to a final concentration of 70% followed by an overnight incubation at −20 °C. The precipitated glycogen was pelleted by centrifugation at 15000 g for 10 min and washed with 70% ethanol and 98% absolute ethanol, consecutively. The precipitated glycogen was dried and digested with 35 U of amyloglucosidase (10115, Sigma Aldrich) in 1 mL of 100 sodium acetate pH 4.5 for 2 h. 200 μl of the samples were mixed with 1 mL of 6% O-toluidine in acetic acid and incubated at 100 °C for 10 min. Absorbance was then read at 635 nm. A glucose calibration curve was used to determine the amount of glycogen in the samples. For every condition, at least three biological replicates were measured.

### Oxygen evolution measurement

Oxygen evolution was measured in vivo using a Clark-type oxygen electrode (Hansatech DW1, King’s Lynn, Norfolk, UK). Light was provided from a high-intensity white light source (Hansatech L2). Oxygen evolution of 2 mL recovering cultures at an OD_750_ of 0.5 was measured at room temperature and 50 μmol photons m^−2^ s^−1^. Three biological replicates per condition were measured.

### Pulse amplification measurement (PAM)

PSII activity was analyzed in vivo with a WATER-PAM chlorophyll fluorometer (Walz GmbH, Effeltrich, Germany). All samples were dark-adapted for 5 min before measurement. The maximal PSII quantum yield (F_v_/F_m_) was determined with the saturation pulse method (Schreiber et al., 1995). Cultures were diluted 1:20 before the measurements in a final volume of 2 mL. Three biological and three technical replicates were measured (three measurements of each biological replicate).

## Supplemental material

**Figure S1.**
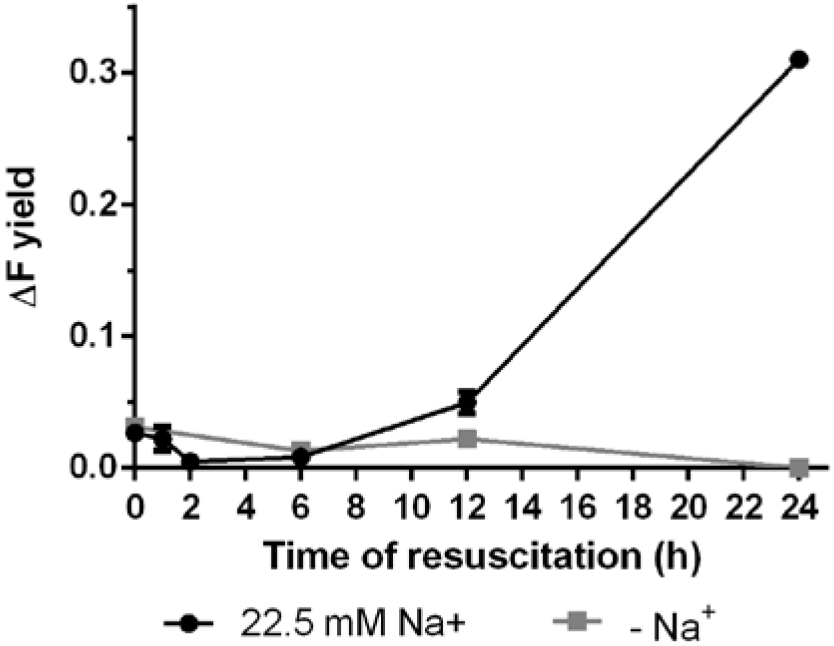
Residual photosynthetic activity is still present during the first hour of resuscitation and it is lost in the absence of sodium. Photosystem II quantum yield determined by pulse-amplitude-modulation (PAM) fluorometry of WT cells during recovery from chlorosis in the presence and absence of sodium. ΔF yield represents the maximal PSII quantum yield (F_v_/F_m_). 3 biological and 3 technical replicates were measured.

**Figure S2.**
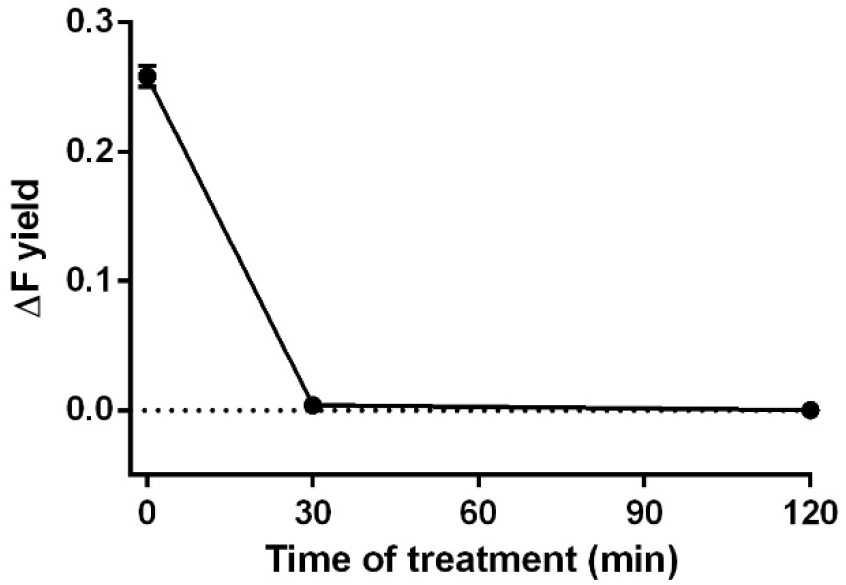
Inhibition of sodium transport blocks photosynthetic activity in vegetative cells. Photosystem II quantum yield determined by pulse-amplitude-modulation (PAM) fluorometry of WT vegetative cells treated with 100 μM EIPA. ΔF yield represents the maximal PSII quantum yield (F_v_/F_m_). 3 biological and 3 technical replicates were measured.

